# Cancer eQTLs can be determined from heterogeneous tumor gene expression data by modeling variation in tumor purity

**DOI:** 10.1101/366922

**Authors:** Paul Geeleher, Aritro Nath, Fan Wang, Zhenyu Zhang, Alvaro N. Barbeira, Jessica Fessler, Robert L. Grossman, Cathal Seoighe, R. Stephanie Huang

**Affiliations:** Section of Genetic Medicine, Department of Medicine, University of Chicago.; Section of Hematology/Oncology, Department of Medicine, University of Chicago.; Department of Experimental and Clinical Pharmacology, University of Minnesota; Ben May Department for Cancer Research, University of Chicago.; Center for Data Intensive Science, University of Chicago; Department of Pathology, University of Chicago, Chicago; School of Mathematics, Statistics and Applied Mathematics, National University of Ireland

**Keywords:** eQTL, GWAS, cancer, gene regulation, deconvolution

## Abstract

Expression quantitative trait loci (eQTLs) identified using tumor gene expression data could affect gene expression in cancer cells, tumor-associated normal cells, or both. Here, we demonstrate a method to identify eQTLs affecting expression in cancer cells by modeling the statistical interaction between genotype and tumor purity. Only one-third of breast cancer risk variants, identified as eQTLs from a conventional analysis, could be confidently attributed to cancer cells. The remaining variants could affect cells of the tumor microenvironment, such as immune cells and fibroblasts. Deconvolution of tumor eQTLs will help determine how inherited polymorphisms influence cancer risk, development, and treatment response.

## Background

Expression quantitative trait loci (eQTLs) have been mapped in many tumor types, including high-profile studies in glioma[1], colon[2], breast[3] and prostate cancer[4]. These studies measured genome-wide gene expression in tumors and identified associations between these gene expression levels and common inherited (germline) genetic variants (e.g. single nucleotide polymorphisms (SNPs)) profiled in the same patients. These results have been very widely applied: For example, the majority of inherited cancer risk variants implicated by GWAS[5] are in non-coding likely-regulatory[6,7] regions of the genome. Thus, to identify genes regulated by these variants, eQTLs identified from tumor tissue[2,3] (and sometimes normal tissue[8]) are typically interrogated—facilitating rational functional follow-up studies[9]. Indeed, inherited genetic variation is associated with the development of specific somatic mutation profiles in cancers, and functional work demonstrated that this can be caused by germline mediated changes in gene expression in cancer cells[10]. Additionally, cancer eQTLs have been extensively studied in the context of pharmacogenomics, for example, inherited variants affect the expression levels of membrane pump/transporter genes modulating chemotherapeutic response[11]. Notably, inherited variants associated with chemotherapeutic response in cell lines are also enriched for eQTLs[12]. Putative drug target genes with existing evidence of disease relevance from genetic association studies are also more likely to be successful in the drug development pipeline; however, this is critically dependent on correctly assigning variants to the genes they regulate[13]. These examples, pertaining to cancer risk, development, and treatment, include only a small subset of applications of cancer eQTL profiles.

However, previous cancer eQTL studies quantified cancer gene expression by extracting RNA from tumor biopsies, which are not a pure sample of cancer cells; instead, these are a heterogeneous mixture of, for example, cancer cells, tumor-infiltrating immune cells, supporting tissue (stroma) and normal epithelial cells from the surrounding tissue. Therefore, the expression profiles obtained reflect both cancer and non-cancer cells. Hence, eQTLs identified this way could arise from cancer cells, tumor-associated normal cells, or both.

Recent studies have developed reliable computational deconvolution methods that use genomics data to estimate the proportion of different cell types in tumor biopsies[14,15], such as those collected by The Cancer Genome Atlas (TCGA). These methods have been shown to accurately recapitulate cell type proportions in controlled experiments, where cell type mixtures are known[16]. Methods have been developed to generate such estimates from gene expression, methylation, and copy number data; these have been compared to estimates from Hematoxylin and Eosin (H&E) staining and it has been observed that all approaches are reasonably concordant, leading to the development of consensus methods, which combine estimates from these approaches[15]. Crucially, these studies have found pervasive differences in tumor purity, both within and across different types of cancer. For example, while samples can be admitted into TCGA with as little as 60% cancer cell content based on H&E staining, the tumor purity inferred from genomics approaches is even lower for some TCGA samples[15]. However, no previous cancer eQTL mapping study has appropriately dealt with the influence of tumor-associated normal cells. In fact, they have essentially treated bulk tumor expression as representative of gene expression in cancer cells. As such, it is plausible that any conclusions drawn about the eQTL landscape of cancer, for example, their similarity to their matched tissue-of-origin[2], could simply result from eQTLs in the tumor-associated normal tissue being misattributed to cancer cells.

In this study, we have developed a statistical approach, which by integrating bulk tumor expression data with estimates of tumor purity, can identify the eQTLs that can be confidently attributed to cancer cells. Using TCGA breast cancer data as a case study and METABRIC as validation, we show that a substantial proportion of reported eQTLs, including known breast cancer risk variants, show no evidence of an effect in cancer cells, but may in fact affect expression in tumor-associated normal cells. Thus, the functional role of these variants must be re-evaluated.

Note: throughout this manuscript we use the terms “bulk tumor” or “tumor” to refer to the heterogeneous mixture of cells found in a solid tumor biopsy; we use “tumor-associated normal” to refer to all non-cancer cell types found in solid tumors (e.g. immune cells, normal epithelial cells) and “cancer” cells to specifically refer to transformed cells.

## Results

### A conventional tumor eQTL mapping strategy will recover eQTLs from both cancer cells and tumor-associated normal cells in simulated data

To establish whether eQTLs in tumor-associated normal cells may indeed influence eQTL profiles recovered from bulk tumor expression data, we first created a simulated dataset where underlying cancer/normal eQTL profiles were known *a priori*. Simulations consisted of expression levels of 600 genes in pure “cancer” samples and in pure “normal” tissue samples. These were then combined to simulate a “bulk tumor” expression dataset, consisting of 1,000 samples. Six classes of eQTLs were created, each represented by 100 genes; these were (1) genes with eQTLs in cancer and normal, but with different effects in the two cell types (2) genes with eQTLs in cancer only (3) genes with eQTLs in normal only (4) genes with no eQTL in either cell type (5) genes with the same eQTL in both cell types and (6) genes with similar eQTLs in both cell types. Because the purpose of these simulations was to study the performance of this model in real cancer data, the parameters, such as sample size, expression levels, effect sizes and proportions of cancer/normal cells, were chosen to resemble the TCGA breast cancer cohort (see Methods).

We applied the current standard eQTL mapping strategy to these simulated data, where the expression levels from bulk tumors were treated as representative of cancer itself (henceforth referred to as the “conventional model”; see Methods). Importantly, the assumption here is that the goal is to identify eQTLs influencing gene expression in cancer cells, therefore true simulated cancer eQTLs were treated as the ground-truth for all statistical measures of performance reported in this and the next section. By comparing the results obtained from the model to the true known cancer eQTLs created as part of the simulation, this approach achieved reasonable sensitivity and specificity (79.5% and 80.3% respectively). However, there was a clear influence of the simulated eQTLs in the normal cells on the recovered effects from bulk tumor expression (Pearson’s correlation (*r*) = 0.9, *P* = 1.3 × 10^−38^ between simulated effect size of eQTLs with an effect in normal but not cancer cells and their estimated effect size from the conventional model; Fig. 1(a)). Furthermore, while we expected a false discovery rate *(FDR;* estimated using the Benjamini and Hochberg approach) of 5%, the true *FDR* was 11.1%, when the known simulated set of cancer eQTLs was treated as the ground-truth. Most (37 of 40) of these false discoveries were falsely attributed associations resulting from eQTLs in normal cells (Supplementary Table 1).

**Figure 1:**
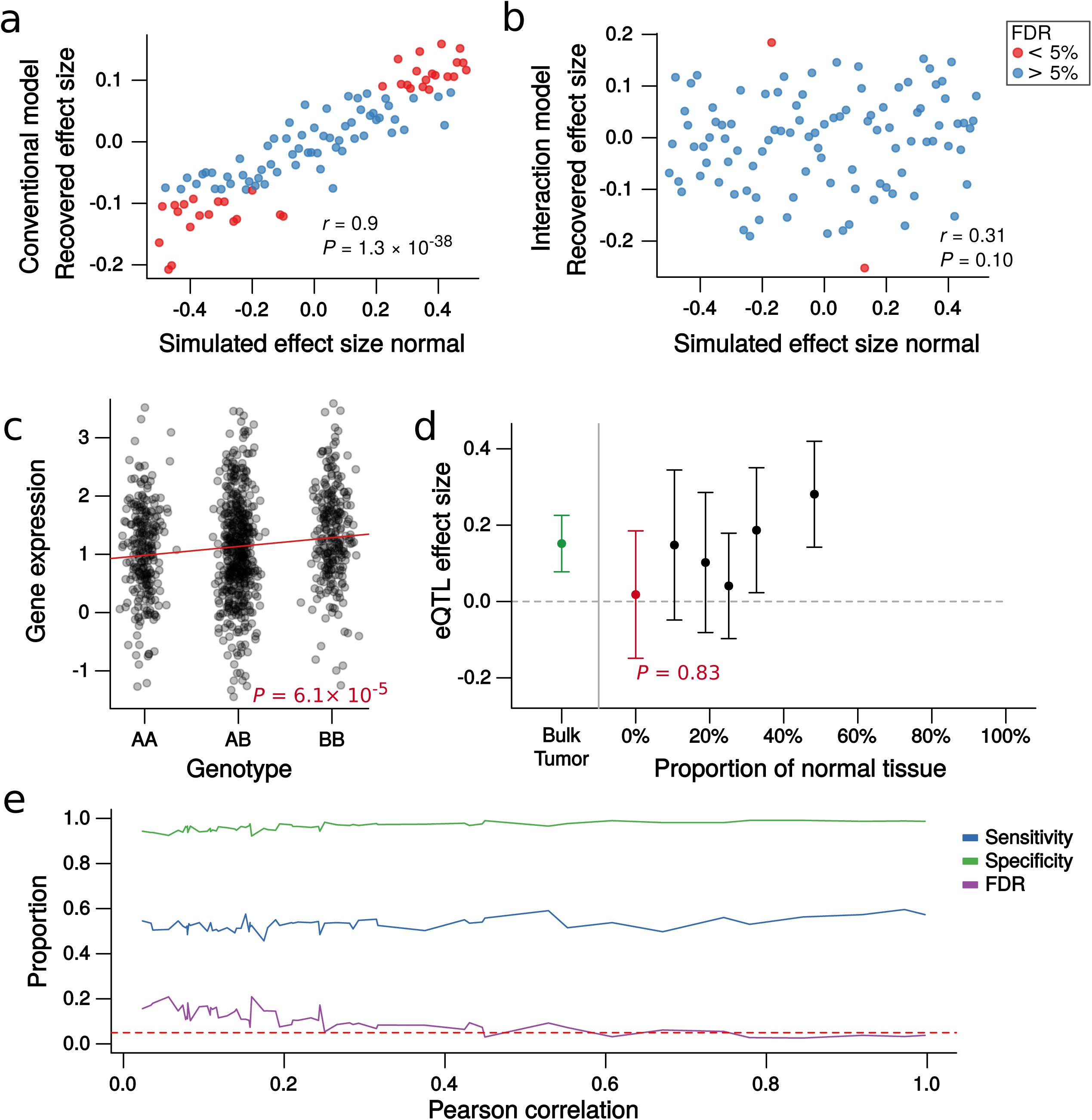
The interaction model can accurately attribute eQTLs to cancer using bulk tumor gene expression in simulated data. (a) Scatterplot of the eQTL effect size recovered from a conventional analysis of bulk tumor expression data (y-axis) against the known normal eQTL effect size created by simulation (x-axis) for the 100 eQTLs that were simulated to have an effect in normal cells, but not cancer. Points are colored red if the conventional model identified these as significant at *FDR* < 0.05. The eQTL effects recovered by the conventional model (y-axis) are heavily influenced by the eQTL effects in tumor-associated normal cells. (b) Scatterplot of the estimated cancer eQTL effect size recovered by the interaction model (y-axis) plotted against the known normal eQTL effect size created by simulation (x-axis) for the same 100 eQTLs as Fig. 1(a) that were simulated to have an effect in normal cells, but not cancer. Points are colored red if the interaction model identified these as significant at *FDR* < 0.05. The recovered eQTL effects (y-axis values) are no longer affected by eQTLs in associated normal cells and in general, have not been misattributed to cancer. (c) Strip chart of a simulated eQTL in tumor expression data, where the effect size in cancer cells was simulated to be 0 (i.e. no eQTL) and the effect size in tumor-associated normal cells was simulated to be 0.48. The conventional model misattributed this eQTL to cancer. (d) The same eQTL as Fig. 1(c), with the effect size calculated in five bins (black points), grouped by the proportion of tumor-associated normal cells. The effect size decreases with increasing proportions of cancer cells. The extrapolated effect size in cancer cells, estimated by the interaction model, is shown in red. The effect size recovered from the bulk tumor, obtained by the conventional model, is shown in green. Whiskers represent 95% confidence intervals. The interaction model has not misattributed this eQTL to cancer cells. (e) The change in the sensitivity, specificity, and *FDR* achieved by the interaction model as the level of noise with which the proportion of cancer cells is measured changes. The ‘Tearson correlation” on the x-axis is the correlation between the known simulated proportions and those “measured” as more noise is added (see Methods). The dashed red line is at 0.05, the rate at which the *FDR* was controlled for these tests using the Benjamini and Hochberg method. The *FDR* is well controlled by the interaction model, even when the correlation between the real and measured (noise added) proportions approaches 0.5. Note: if the cancer cell proportions are completely randomized, the true *FDR* is 22% (at the 5% threshold). Again, when calculating these true *FDRs*, the known simulated set of cancer eQTLs were treated as the ground truth.

### Cancer eQTLs can be accurately identified from bulk tumor expression data by modeling the interaction of tumor purity and genotype in simulated data

To recover cancer eQTLs from bulk tumor expression data, we have built upon (see Methods) a previous study to identify eQTLs with different effects in human neutrophils and lymphocytes using whole blood expression data[17]. Like conventional eQTL mapping, our new approach involves fitting a linear regression model of gene expression level against genotype: However, in addition to genotype, the estimated proportion of tumor-associated normal cells (tumor purity) is included as a covariate, as well as the interaction between the estimated tumor purity and genotype (henceforth referred to as the “interaction model”; see Methods). Critically, the estimate of the main effect associated with this interaction term allows the eQTL to be assigned to cancer, not the interaction term itself (see Methods). Intuitively, this works by estimating how the magnitude of the association between bulk tumor gene expression and genotype changes as a function of the proportion of cancer/normal cells, then extrapolating the effect size to 100% cancer cells. Under reasonable assumptions, we have proved this approach mathematically and demonstrated how this model should be interpreted (see Supplementary Information — Model Derivation).

The interaction model recovered simulated cancer eQTLs with a sensitivity and specificity of 58.3% and 96.1% respectively. A small drop in power (Supplementary Tables 1 & 2; Supplementary Figure 1) was expected given the extrapolation to a cell type-specific state and the simulations taking account of the potential for shared eQTLs between cancer and normal cell types. However, the true *FDR* dropped to 3.3%, below the expected rate of 5%. Only two “normal only” (group 3; see Methods) eQTLs were misattributed to cancer and the influence of normal cells observed for the conventional model was eliminated (Fig. 1(b); Supplementary Table 2). To further illustrate the utility of the model, a normal-driven eQTL analyzed with a conventional model is shown in Fig. 1(c), along with the capacity of the interaction model to extrapolate the correct effect size in cancer cells, deducing that this signal was driven by samples with large quantities of tumor-associated normal cells (Fig. 1(d)).

In cancer eQTL mapping, the assumption has been implicit that the eQTLs identified from tumor samples affect gene expression in cancer cells. However, the pervasive genomic aberrations and dysregulation of key master regulators that occurs in cancer cells[18] could obscure or eliminate associations between germline polymorphisms and gene expression, either by increasing transcriptional noise or by disrupting the regulatory landscape. Thus, the inherited genetic influence on gene expression could be far greater in normal cells than in cells that have undergone neoplastic transformation. To assess the plausibility that eQTLs previously discovered from tumor expression data could be largely driven by normal cells, we included an additional 500 genes with “normal only” eQTLs in our simulated dataset. Again, assuming the objective is to identify eQTLs that affect gene expression in cancer cells, a conventional model applied to bulk tumor expression data performs very poorly. Using an *FDR* threshold of 5% we in fact observed a rate of false discovery rising to 46% of significant associations (Supplementary Table 3). Of the 270 false discoveries, 267 were misattributed eQTLs affecting gene expression in normal cells only. However, when the interaction model was used, the rate of false discovery was again accurately controlled (3% false discoveries at an imposed *FDR* threshold of 5%) and only 5 eQTLs in normal cells (<1%) were misattributed to cancer. Furthermore, the interaction model could accurately identify true cancer eQTLs even when tumor purity was measured with noise similar to levels expected in real data[19] (Fig. 1(e); see Methods for details). Notably, just including the proportion of cancer cells as a covariate in a conventional model had no impact on the performance, with the observed *FDR* remaining at 45.9% (at the imposed 5% threshold; Supplementary Table 3). Thus, tumor purity cannot simply be “accounted for” by including it as a model covariate or including surrogate variables that approximate tumor purity such as principal components or PEER factors—modeling the interaction of tumor purity and genotype is absolutely critical to correctly assign eQTLs to cancer cells. Ignoring this can potentially falsely attribute enormous numbers of eQTLs from tumor-associated normal cells. Notably, simply restricting to tumors with higher cancer cell content is also likely not an optimal solution this problem; doing so caused a large drop in sensitivity compared to the interaction model, at a true *FDR* < 5% (Supplementary Figure 2).

While no simulated dataset can capture the full complexity of *in vivo* biology, these analyses suggest that (i) it is plausible that many, if not most eQTLs identified from tumor expression data using conventional approaches actually affect gene expression in normal cells, not in cancer cells and (ii) using the parameters of the TCGA breast cancer data, modeling the interaction of tumor purity and genotype performs well at correctly attributing true cancer eQTLs. Below, we perform a case-study using an integrative analysis of real data from TCGA breast cancer, breast cancer GWAS results, and samples from the Genotype-tissue Expression (GTEx) project.

### Case-study: mapping *cis*-eQTLs in breast cancer

To test the utility of the interaction model on real data, we conducted *cis*-eQTL mapping in TCGA breast cancer samples, where both germline genotype and bulk tumor RNA-Seq data were available (n = 894). We also applied a conventional model to bulk tumor expression data (see Methods). We focused on breast cancer as it has the largest available sample size, and is reasonably representative of tumor types with high normal cell contamination (Fig. 2(a)). We estimated tumor purity using a consensus approach that combined the estimates from copy number variation, gene expression, DNA methylation and H&E staining[15]. Tumor purity varied substantially in TCGA breast cancer samples (Fig. 2(b)) and was significantly correlated with the expression of 11,927 of 15,574 genes *(FDR* < 0.05; Fig. 2(c)), highlighting the obvious potential of eQTLs in these normal cells to influence eQTL profiles inferred from bulk tumor expression.

**Figure 2:**
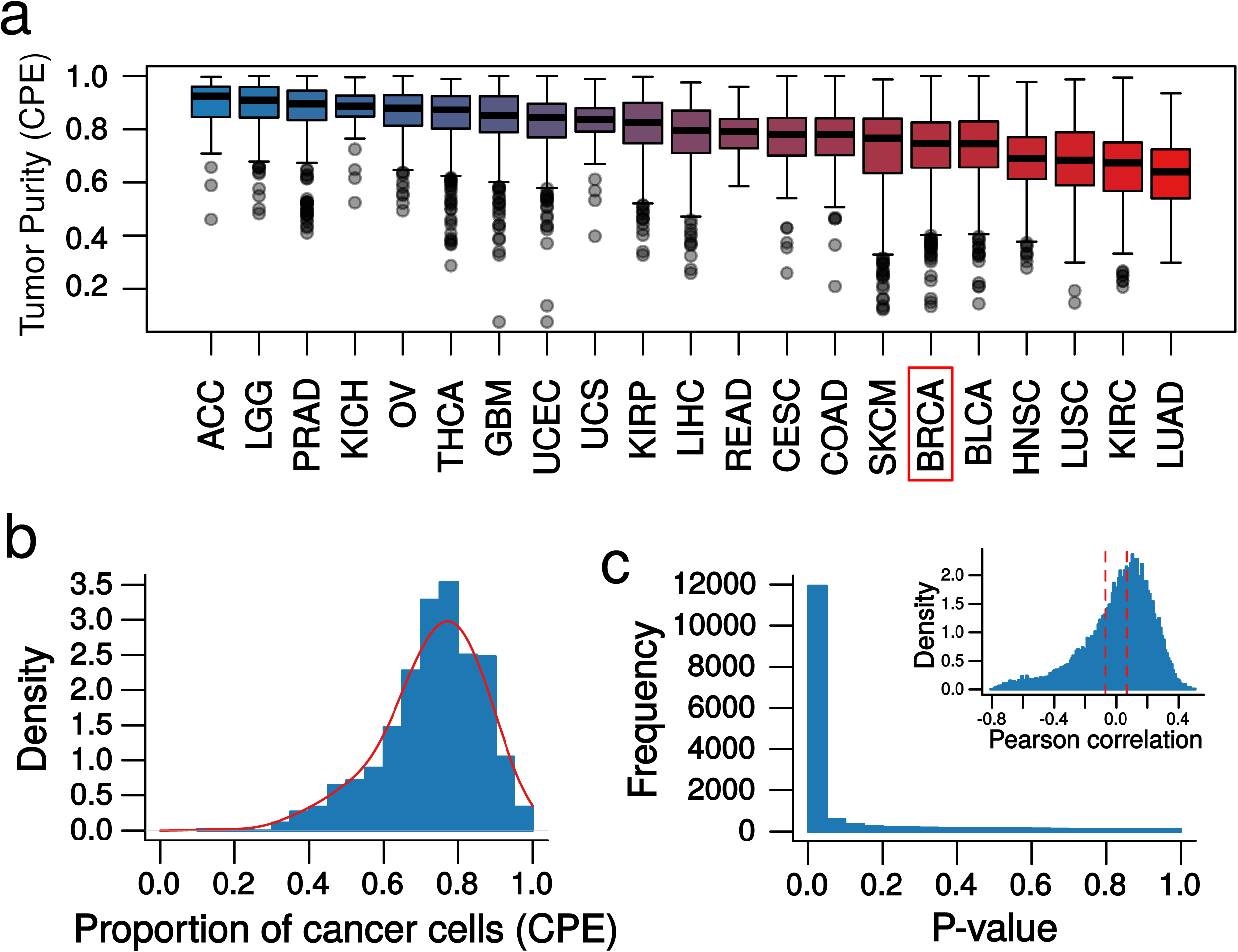
Estimates of tumor purity in TCGA samples vary substantially with an between cancer types. (a) Boxplot of tumor purity estimates from Consensus Purity Estimation (CPE) [15] method for 21 solid tumor types in TCGA. Breast cancer is highlighted in red. (b) Histogram (blue) and density plot (red) of variability in tumor purity for the TCGA breast cancer samples (n = 1,063), estimated using CPE. (c) Histogram of *P*-values for the association of expression and tumor purity; 76.5% of genes’ expression were significantly correlated with tumor purity *(FDR* < 0.05). Corresponding Pearson’s correlation values of gene expression and CPE estimates of tumor purity in TCGA breast tumors are shown in the inset. The area outside the red dashed lines represents significant correlations *(FDR* < 0.05).

We evaluated 3,602,220 associations between tag SNPs and the expression levels of genes within 500 kilobases of each tag SNP. The data were filtered and preprocessed based on the recent guidelines of GTEx, including steps to control for population structure, unmeasured confounders, and expression heterogeneity (see Methods). We identified 57,189 significant *cis*-eQTL associations *(FDR* < 0.05; Fig. 3(a)) using the conventional model. However, using the interaction model, just 8,833 eQTLs could be confidently attributed to cancer cells *(FDR* < 0.05; Fig. 3(a)). Of the 8,833 associations attributed to cancer cells, 7,542 were also identified by the conventional model and 751 were novel. Results were similar when copy number or methylation were included as an additional covariate (as per Li *et al*.[3] (Supplementary Figure 3)) and when samples were grouped by subtype (Supplementary Tables 4 & 5; see Methods). When we randomly permuted the tumor purity estimates, the number of eQTLs that could be attributed to cancer cells was just 239 (Fig. 3(a)). We show a specific example in Figs. 3(b-e) to illustrate the process of attributing eQTLs to the affected cell type. In this example, the association between SNP rs6458012 and the expression of *MDGA1* in breast tumors (*P* = 1.5 × 10^−29^; Fig. 3(b)) could not be attributed to breast cancer cells (*P* = 0.26; Fig. 3(c)), given the tumor purity values. This eQTL likely arises from tumor-associated stromal and immune cells, in which the genotype of this locus has a strong effect on gene expression with the same directionality as the eQTL estimated by the conventional model (*P* = 6.3 × 10^−13^ and 1.4 × 10^−10^ in GTEx transformed fibroblasts and lymphoblastoid cell lines (LCLs), respectively; Fig. 3 (d & e))

**Figure 3:**
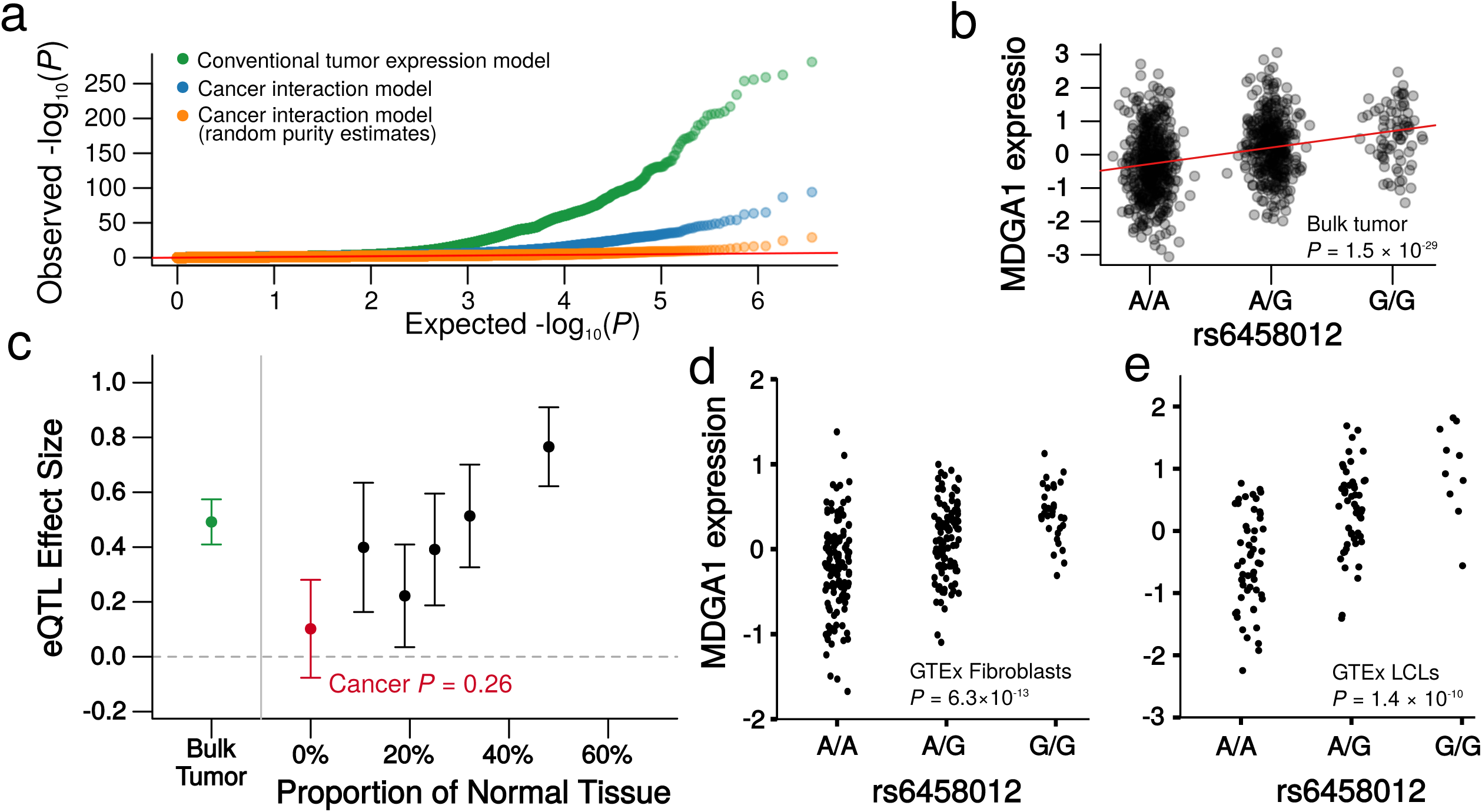
Using the interaction model to identify cancer eQTL profiles from TCGA breast bulk tumor gene expression data. a) QQ-plot of P-values for *cis*-eQTL associations in TCGA breast cancer samples, recovered using a conventional eQTL analysis of bulk tumor gene expression data (green) when eQTLs are attributed to cancer using the interaction model (blue), and when tumor purity estimates in the interaction model are randomly permuted (orange). Observed *P*-values (y-axis) are plotted against the uniform distribution of *P*-values (x-axis). b) A strip chart showing the association of rs6458012 and the expression *MDGA1* in TCGA breast cancer tumors, with the association identified by the conventional model shown as a red line. c) Plot deconstructing the association between rs6458012 and *MDGA1*. Points are effect sizes and whiskers represent 95% confidence intervals. The association from the conventional model applied to TCGA breast cancer bulk tumors is shown in green (corresponding to Fig. 3(b)). Shown in black are the effect sizes and confidence intervals for the association of rs6458012 and *MDGA1* when TCGA breast samples are divided into five equally sized bins, based on each sample’s estimated proportion of cancer cells. The estimated effect size decreases as the proportion of cancer cells decreases. The extrapolated effect size in cancer cells, estimated by the interaction model, is shown in red; this association is not statistically significant, illustrated by the 95% confidence interval overlapping the grey dashed line, which represents an effect size of 0. This suggests the association recovered by the conventional model did not arise in cancer cells. d) A strip chart showing the association of rs6458012 and the expression *MDGA1* in fibroblasts from GTEx. These are associated (*P* = 6.3 × 10^−13^) with the same directionality as identified in TCGA breast tumors (Fig. 3(b)). e) A strip chart showing the association of rs6458012 and the expression *MDGA1* in LCLs from GTEx. These are associated (*P* = 1.4 × 10^−10^) with the same directionality as identified in TCGA breast tumors (Fig. 3(b)).

### The interaction model attributes fewer immune and fibroblast-specific eQTLs to breast cancer cells in the TCGA cohort

As outlined above, when the interaction model was used, we found that the majority (49,647; 86.8%) of the eQTLs identified from bulk tumor expression data could not be attributed to cancer cells. Indeed, 18,595 of these potentially falsely-attributed eQTLs were also eQTLs, with concordant directionality in one or more of normal breast (8,536 eQTLs), LCL (4,531 eQTLs) or fibroblast (15,810 eQTLs) tissues in GTEx. However, cancer eQTL profiles have never been studied in the absence of normal cells and germline genotypes are not typically collected from cell line donors; hence, there is no established gold standard to compare the sensitivity/specificity of the conventional and interaction models in real data. However, we can assess whether the interaction model eliminates associations for likely immune and stromal cell-specific eQTLs. To do this, we used GTEx data to define a set of eQTLs that were likely to be misattributed; i.e. they were *more likely* to have arisen in immune and stromal cells, rather than from breast cancer cells. We defined this set as *cis*-eQTLs identified in lymphoblastoid cell lines (LCLs) or transformed fibroblasts in GTEx *(FDR* < 0.05), which were not even nominally significant (P > 0.05) in GTEx breast tissue. We reasoned that LCLs and fibroblasts provide a good proxy for tumor-associated immune and stromal cells, while the regulatory landscape of breast cancer cells is likely to maintain a similarity to breast, the tissue from which they developed. These criteria yielded a set of 47,196 eQTLs shared between GTEx and TCGA that had a higher likelihood of being misattributed if identified as cancer eQTLs. Of the 57,189 significant associations from the conventional model, 5,440 were among this set defined as likely arising in associated-normal cells. For 8,833 associations from the interaction model, this number was reduced to 572. This is a significant reduction in the proportion of these likely misattributed eQTLs (Fig. 4(a & b), *P* = 8.1 × 10^−22^ from Fisher’s exact test, odds ratio 1.51). Thus, consistent with our simulations, there is convincing evidence in real data that the use of the interaction model reduces the misattribution of eQTLs from tumor-associated normal cells. Furthermore, we also mapped breast cancer eQTLs using only 10% of the TCGA breast cancer samples that had the highest estimated cancer cell content (all > 88.6% purity; median = 91.2%, n = 89). As expected, the eQTL effects estimated from this high purity subset were (globally) much more similar to those estimated by the interaction model compared to the conventional model (r = 0.447, 95% CI = 0.446-0.448 for the interaction model; *r* = 0.299, 95% CI = 0.299-0.3 for the conventional model; Supplementary Figure 4).

**Figure 4:**
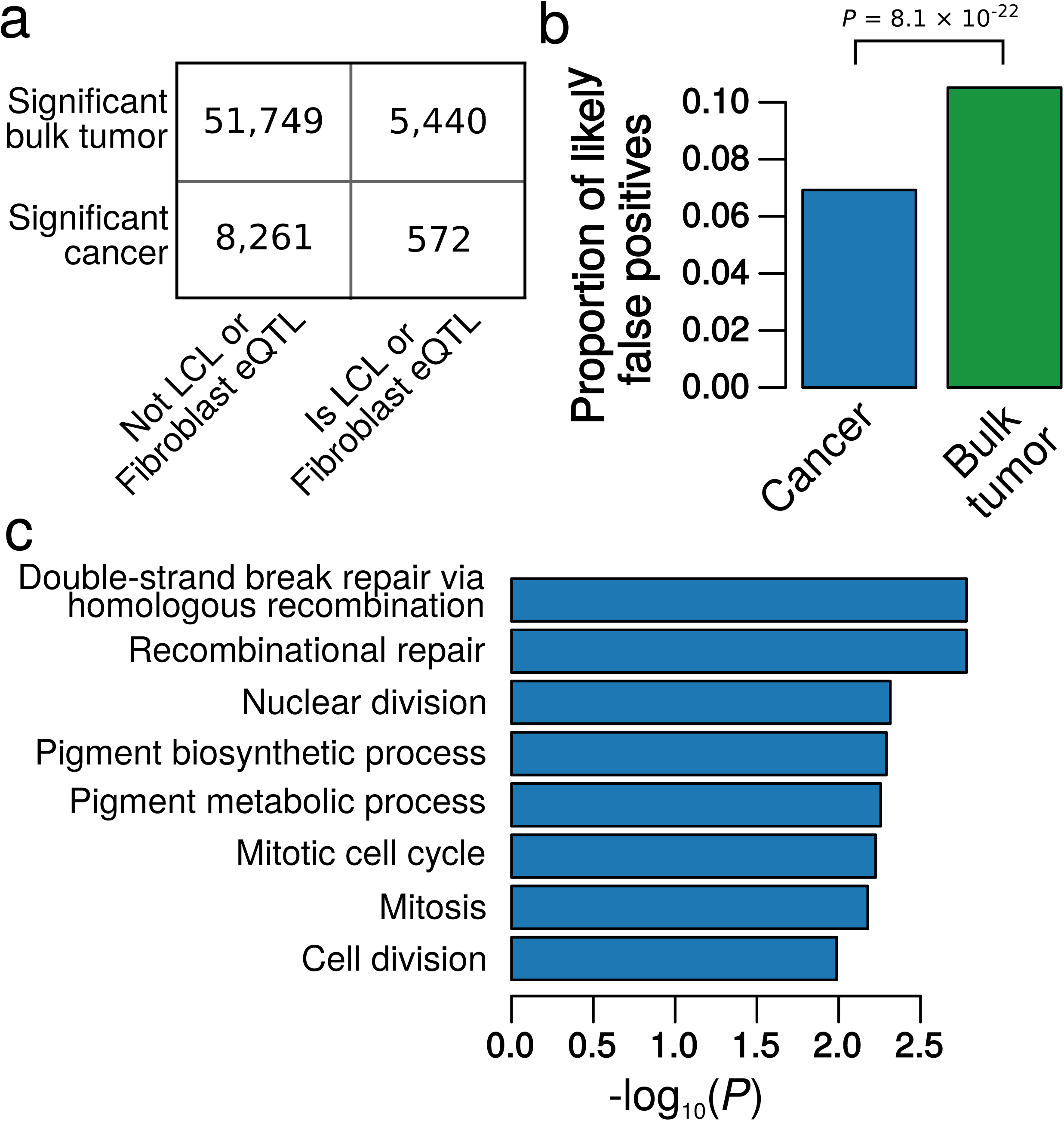
Classification and annotations for eQTLs identified using a conventional or an interaction model. (a) 2 × 2 contingency table showing the number of total eQTLs identified using either the conventional (tumor) model, or the interaction (cancer) model, which were also significant *(FDR* < 0.05) in GTEx LCL or fibroblast data, but not GTEx breast data (*P* > 0.05). (b) The proportions of eQTLs identified by the conventional (bulk tumor) or interaction (cancer) model that are likely to be derived from normal cells, based on Fig. 4(a). (c) Bar graph of the lowest 8 *P*-values of 3,679 GO biological processes tested; *P*-values are for the enrichment of genes whose eQTL profile changes between TCGA breast cancer (identified by the interaction model) and normal breast tissue in GTEx.

### eQTLs that are disrupted following tumorigenesis tend to affect genes involved in cancer-relevant processes

We also expect that genes whose regulation is disrupted following tumorigenesis would be more likely to be involved in cancer hallmark processes[20,21]. Thus, for all *cis*-eQTLs represented in GTEx breast tissue and TCGA breast cancer, we compared the magnitude of the effect of each eQTL between the two datasets (see Methods). For 3,885 of 3,270,829 eQTLs, there was evidence *(FDR* < 0.05; Supplementary Figure 5; Supplementary Table 6) of a difference between breast cancer and normal breast tissue. Of these, 3,068 had a larger effect (comparing absolute values) in normal breast tissue and 797 in cancer. We compiled a list of eQTL-associated genes for which there was evidence of a difference in this germline mediated regulation of gene expression between cancer and normal cells. Then, to determine whether these changes were biologically meaningful we assessed these genes for enrichment of Gene Ontology (GO) biological process (see Methods). Indeed, the most strongly enriched processes included cancer relevant terms (Fig. 4(c); Supplementary Table 7; Supplementary Figures 6 & 7). The top associations included DNA repair and cell cycle, key process influencing breast cancer susceptibility and progression. Some of this dysregulation may be attributable to increased expression heterogeneity or different expression levels among these genes in cancer and understanding the mechanisms by which normal regulation of these genes is disrupted will represent a starting point for future mechanistic studies.

### Validation of TCGA breast cancer findings in the METABRIC dataset

Next, we sought to replicate our results using an additional 997 breast tumor expression profiles and genotypes generated by the METABRIC consortium[22]. Although this is the most suitable validation cohort available, there are some limitations to this dataset; for example, the genotypes were generated from (less reliable) tumor tissue (see Methods), and expression was estimated using a microarray platform, which is likely less sensitive than the RNA-seq platform used by TCGA. Despite this, the results were similar to TCGA. Using a conventional model 47,354 eQTLs were identified *(FDR* < 0.05) in METABRIC and this number dropped to 9,235 when the interaction model was applied, with an overlap of 8,142. Thus, similarly to the TCGA cohort, most tumor eQTLs identified in METABRIC could not be confidently attributed to cancer cells. Despite the differences between these datasets, the overlap of eQTLs identified in TCGA and METABRIC was much higher than expected by chance for both the conventional and interaction models: 39.4% of tumor eQTLs identified *(FDR* < 0.05) by the conventional model in TCGA were also significant *(FDR* < 0.05) when the conventional model was applied to METABRIC (57.4% reached *P* < 0.05). 31.5% of cancer eQTLs identified *(FDR* < 0.05) by the interaction model in TCGA were also significant *(FDR* < 0.05) when the interaction model was applied to METABRIC (52.4% reached *P* < 0.05). A slight drop in this replication rate for the interaction model was expected given the additional challenge of assigning eQTLs to a specific cell type, rather than just identifying bulk tissue eQTLs.

### Correctly assigning bulk tumor eQTLs can inform the biological consequences of breast cancer risk variants identified by GWAS

GWAS have revealed many common genetic variants associated with cancer, including high-profile studies of breast cancer risk[6,23]. eQTL mapping represents an important early step in characterizing the function of cancer risk variants, most of which lie outside protein-coding regions[3,24,25]. Thus, we re-analyzed the eQTL profiles of the variants identified by a recent meta-analysis of GWAS data for breast cancer risk, which identified over 90 loci[6]. 24 of 565 possible SNP-gene *cis*-eQTL pairs were significant *(FDR* < 0.05) when a conventional model was applied to the TCGA breast tumor expression data (arising from 16 of the 81 risk SNPs that could be mapped to one-or-more genes; see Methods). However, 9 of these eQTLs were not even nominally significant (P > 0.05) when extrapolated to cancer cells using the interaction model, suggesting these are strong candidates for eQTLs arising from normal cells. Indeed, all of these 9 associations were significant in at least one of fibroblast, breast or LCLs in GTEx, in all cases with the same directionality as the eQTL effect estimated from bulk tumor expression using the conventional model (Fig. 5(a & b); Supplementary Table 8).

**Figure 5:**
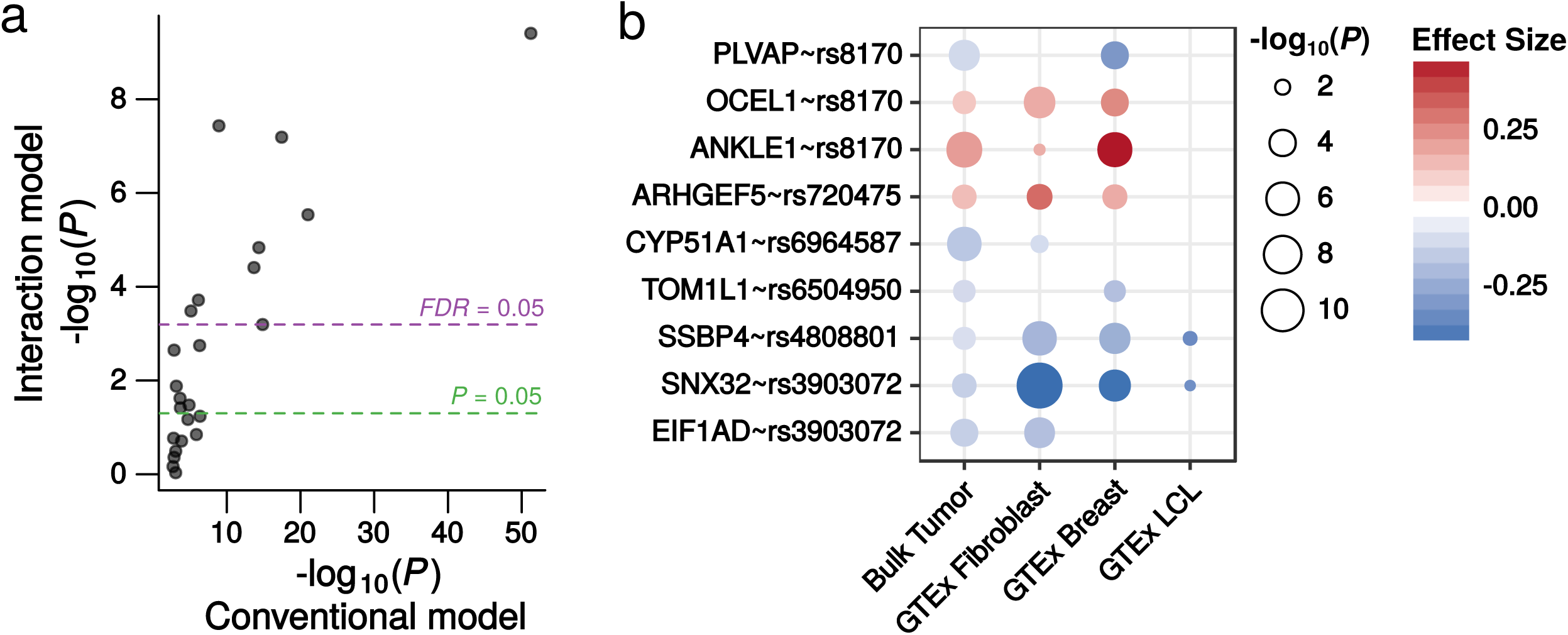
Conventional model eQTLs for breast cancer risk GWAS significant genetic variants, which cannot be attributed to cancer cells by the interaction model, are all statistically significant with concordant directionality in GTEx LCLs, breast or fibroblasts. (a) Scatterplot of eQTLs identified for breast cancer risk variants from a meta-analysis of GWAS data[6]. −log_10_ *P*-values for significant eQTLs *(FDR* < 0.05) using a conventional model in TCGA breast cancer tumors are shown on the x-axis. –log_10_ *P*-values from the interaction model for these same eQTLs are shown on the y-axis. Significance thresholds of 0.05 are shown for the interaction model (green and purple dashed lines). 15 of these 24 eQTLs were no longer significant *(FDR* > 0.05) when the possibility of these eQTLs arising from tumor-associated normal cells is modeled. (b) P-values and effect sizes for the nine eQTLs in Fig.5(a) that were no longer even nominally significant (*P* > 0.05) when the interaction model was used. For each eQTL, the effect size is represented by the red-blue divergent color scale and the *P*-value by the size of the point; these are shown for TCGA bulk breast tumor (i.e. the conventional model) and transformed fibroblast, breast tissue and LCLs from GTEx. All of these eQTLs are evident in at least one of these GTEx tissues, with 100% concordant direction of effect as identified from TCGA breast bulk tumor expression using the conventional model. Points are shown for associations with *P* < 0.05.

Using the interaction model, another 9 of these 24 SNPs could be confidently assigned to cancer cells *(FDR* < 0.05; Supplementary Figure 8). Five of these were strong cross-tissue eQTLs in GTEx (for *ATG10, ATP6AP1L* and *RPS23*, all associated with rs7707921, and *C5orf35 (a.k.a. SETD9)* & rs889312; *P* < 1 × 10^−5^ in at least 19 tissues with concordant directionality) and maintain their regulatory capacity in breast cancer cells. Interestingly, 8 of the 9 eQTLs for these GWAS variants, which could be confidently attributed to breast cancer cells, were also at least borderline significant in normal breast tissue in GTEx (1.3 × 10^−20^ < *P* < 7.4 × 10^−2^; Supplementary Figure 8; Supplementary Table 8), suggesting that the effect of genetic variation on gene expression in the baseline normal tissue state is generally maintained following tumorigenesis. However, there is an exception for the SNP rs204247, which affected the expression of *RANBP9* in breast cancer cells only. *RANBP9* is ubiquitously and highly expressed in human tissues (Supplementary Figure 9) and breast cancer cell lines[26] (Supplementary Figure 10), but this eQTL is only evident in esophagus mucosa[27] (*P* = 2 × 10^−8^) and aorta (*P* = 3.9 × 10^−5^) in GTEx (Supplementary Figure 11). rs204247 tags the promotor of *RANBP9*, as well as upstream putative enhancers (in MCF7 breast cancer cells; Supplementary Figure 12). The interaction model indicates that the risk allele (G; per-allele odds ratio = 1.06 (95% CI = 1.03 - 1.1)) increases the expression of *RANBP9* in breast cancer cells (Supplementary Figure 13). Consistent with an oncogenic effect, *RANBP9* is overexpressed in a similar proportion of breast cancer patients as *ERBB2*—an important driver of breast cancer (13.04% and 14.13% respectively[28]). Amplifications of *RANBP9* occur in breast cancer *in vivo* (Supplementary Figure 14) and are associated with increased gene expression (Supplementary Figure 15); although amplifications are less common than for *ERBB2*, suggesting other mechanisms more typically driving its overexpression. Given that *RANBP9* is ubiquitously expressed, this eQTL in cancer cells cannot be explained by the activation of the gene and must reflect some change in gene regulation. Thus, the cancer cell eQTL analysis suggests *RANBP9* may be an important driver of breast cancer risk and progression and the possible oncogenic effects of this gene could represent an interesting starting point for functional studies.

## Discussion

We have demonstrated an improved eQTL mapping strategy for cancer, which uses tumor purity estimates and bulk tumor gene expression data to identify eQTLs that can be confidently attributed to cancer cells. In breast cancer, the result is that most bulk tumor eQTLs cannot be confidently attributed to cancer cells, once the possibility of these eQTLs arising from tumor-associated normal cells is appropriately modeled.

We demonstrated the implications for the interpretation of genetic variants associated with cancer risk. The mechanism-of-action of most cancer GWAS variants remains unknown. However, if these variants affect gene expression in tumor-associated normal cells, but not cancer or baseline normal cells, their disease relevance could lie in modulating how the host—and in particular the cells of the tumor microenvironment—responds to the disease rather than reflecting functions intrinsic to cancer (or pre-cancer) cells themselves. Furthermore, we also showed that one breast cancer risk variant, rs204247, is an eQTL for *RANBP9* in breast cancer cells, but not tumor-associated normal cells. If rs204247 affects *RANBP9* expression only in breast cancer cells, and this is indeed the mechanism by which this SNP pre-disposes individuals to cancer, then some earlier aberration, for example, the activation of a transcription factor, must be a pre-requisite for rs204247’s pathogenic effect. Such an aberration might occur in pre-cancer cells, with individuals carrying the risk allele of rs204247 then manifesting the oncogenic effects of increased *RANBP9* expression. Interestingly, *RANBP9* has been shown to interact with oncogene c-MET, a key regulator in development and cancer stem cells. This interaction has been shown to stimulate RAS signaling, which is crucial to cancer-relevant processes such as cell differentiation, apoptosis, and motility[29], thus offering a possible oncogenic mechanism of this GWAS risk allele. Notably, if this hypothesis is correct, rs204247 is likely affecting druggable pathways. However, this association would not have been apparent by only interrogating baseline normal tissue(s).

In the future, one approach to cancer eQTL mapping will likely be to apply single-cell gene expression methods to tumors—directly measuring gene expression in cancer and tumor-associated normal cells. For many cancer types this should be possible, but currently, single-cell expression datasets are not on a scale required to map eQTL profiles. For the foreseeable future, sample sizes available for gene expression in bulk tumors will remain orders-of-magnitude larger than single-cell datasets. Furthermore, single-cell methods bring additional biases, for example isolating single cells can cause marked changes in expression and low starting amounts of RNA leads to high levels of technical variability[30]. These studies have also encountered difficulty in isolating some cell types from tumors[18]. Hence, mapping the genetic determinants of gene expression in cancer cells, using expression data from bulk tumors, will complement any single-cell studies conducted should the technology become sufficiently well developed and low-cost that it becomes feasible on a suitably large scale. Notably, one immediate benefit of single-cell datasets may be improved signatures to estimate cell type proportions from bulk tumor data.

Here, we have treated breast tumors as composed of two broad cell types, cancer and normal. Of course, these cell types can be further subdivided. The normal component is composed of endothelial, epithelial, stromal and immune cells, which can themselves be subdivided. Cancer cells are also heterogeneous—for example, the presence of stem-like cells. However, differentiating between the eQTL profiles of every cell type would require an interaction term for each cell type. One would also need to be sufficiently confident that the cell type proportions were being accurately estimated, which becomes more difficult given more similar expression profiles in less distinct subtypes. Single-cell gene expression analyses of breast cancer have already shown that cancer and normal cells strongly cluster in principal component analysis[18], meaning breast cancer cells are transcriptionally much more similar to each other than they are to tumor-associated normal cells. Thus, our approach provides a mechanism to identify eQTLs that can be confidently attributed (wholly or in part) to cancer cells from tumor expression data. However, future research in the development of statistical methods for analysis of tumor expression, or single cell-based analyses, could benefit from further interrogating these complexities.

Another assumption that our model makes, is that the presence/absence of normal cells does not itself affect eQTLs in cancer cells, which could result in normal cells influencing tumor eQTL effect-sizes in a non-linear fashion. While previous studies have shown that this linearity assumption is reasonable for expression data[19], for genes where this is not true, it may be difficult or impossible to separate the eQTL profiles of tumor-resident cancer and normal cells using any method, including single-cell RNA-seq.

Additionally, our model, or any such model, cannot prove a non-association. It is incorrect to conclude that tumor eQTLs that cannot be attributed to cancer cells are definitely not eQTLs in cancer cells, or are certainly eQTLs in tumor-associated normal cells. The correct interpretation is that there is no statistical evidence for this eQTL in cancer cells at the current sample size and given factors such as the accuracy with which the data were measured. Notably, cancer eQTLs identified by the interaction model may still be eQTLs in other tumor-associated normal cell types and these should not be interpreted as exclusively-cancer eQTLs.

## Conclusion

We have elucidated a major shortcoming of current eQTL mapping strategies in cancer, in that eQTLs identified from tumor expression data could arise from either cancer or tumor-associated normal cells. We have also proposed a solution, which allows us to recover eQTL profiles for constituent cell types using expression data collected in a mixture of cell types. We have applied this solution to breast cancer, where we showed that most eQTLs discovered in tumors cannot be confidently attributed to cancer cells, once the possibility of these signals arising in tumor-associated normal cells is appropriately modeled. Overall, this work will improve the understanding of gene regulation in cancer, including studying inherited cancer risk variants, disease development, and drug response. This study also provides improved theoretical groundwork for deconvolution of eQTLs effects in other mixtures of cell types, including normal human tissues.

## Methods

### Simulating bulk tumor expression data as a product of underlying “cancer” and “normal” expression data

We simulated cancer and normal gene expression datasets for 600 genes in 1000 samples—the approximate number of patients in the TCGA breast cancer dataset. Cancer and normal expression datasets were then combined to create a bulk tumor expression dataset, with each gene combined using a weighted mean based on purity estimates for the sample. Combining expression datasets in this way assumes a linear relationship between expression levels in the pure and mixed samples, which has previously been shown to be reasonable[19]. For all simulated SNPs, the two alleles were simulated as occurring at an equal frequency (i.e. 500 homozygotes and 250 of each heterozygous group, one of which was arbitrarily designated the minor allele). Simulated eQTL effect sizes (the fold-change in gene expression with each copy of the minor allele) were drawn from a uniform distribution, which ranged from −.5 to .5, in steps of 0.01; this range was chosen as it covers the approximate range of the effect sizes observed in the TCGA breast cancer data. Before adding eQTL effects, the expression level of each allele was randomly sampled from a normal distribution of mean 1 and standard deviation 1 (TCGA expression data were also mapped to a normal distribution of standard deviation 1 (see below)). The 600 simulated genes were split into 6 groups of 100, each of which was treated differently, to represent the likely different types of scenarios that may arise *in vivo*: In Group 1, eQTL effects were introduced in both cancer and normal expression datasets, but the effects were randomly shuffled across genes, representing a scenario where there is an independent eQTL effect on each gene in both cancer and normal tissue. In Group 2, eQTL effects were only introduced in the cancer expression data. In Group 3 eQTL effects were only introduced in the normal expression data. In Group 4 eQTL effects were not introduced in either. For genes in Group 5, the same eQTL effect was introduced in both expression datasets. In Group 6, eQTL effects were simulated to be similar in cancer and normal tissues, by simulating identical eQTLs then adding randomly generated noise in the normal expression data.

Simulated purity estimates were derived from 1,000 randomly chosen consensus purity estimates[15] in real TCGA breast cancer samples. When recovering the cancer eQTLs using the interaction model, noise was added to the purity estimates, to simulate the fact that in real data these estimates will be imprecise: For each sample, noise was added by randomly sampling a normal distribution with mean 0 and standard deviation 0.1; the resulting values were then quantile normalized to the original purity estimates, thus preserving the distribution of the data precisely (Supplementary Figure 16). For Fig. 1(e), the standard deviation of the noise generating normal distribution was varied from 0.01 to 1.5 in steps of 0.025, thus simulating the effects of varying levels of error in the tumor purity estimates; the resulting vector was quantile normalized to the original vector and the Pearson’s correlation shown on the x-axis of Fig. 1(e) were calculated between this noise-added vector and the original vector. All simulations were performed in R and code to reproduce them is available in our supplementary materials.

### Data processing and eQTL mapping in TCGA breast cancer samples

RNA-Seq data for TCGA breast cancer samples were obtained from FireBrowse and filtered to only include primary tumor expression data. These data had been summarized to gene level transcript per million (TPM) estimates using the RSEM[31] software. Corresponding genotype calls, which had been generated using Affymetrix Genome-Wide Human SNP Array 6.0 on blood samples, were obtained from the Genomics Data Commons (GDC)[32]— permission was obtained to download these data. Processed methylation data and copy number data were also obtained from FireBrowse; gene level copy number was estimated as previously described [33]. PAM50 subtypes were obtained from the supplementary materials of Netanely *et al*.[34]. Gene expression and genotype data were pre-processed and filtered primarily using the guidelines of GTEx: Expression data were quantile normalized. The expression of each gene was then mapped to a standard normal distribution, with mean 0 and standard deviation of 1. Genes not expressed in at least 75% of samples were removed. SNPs with a minor allele frequency (MAF) of less than 5% were removed. Males, as well as Y chromosome SNPs and genes, were removed. We estimated ancestry using the first 3 principal component of the genotype matrix. To account for expression heterogeneity and unmeasured confounders, we also estimated 35 PEER factors[35]. The filtering steps yielded 15,574 genes and 701,700 SNPs in 894 breast cancer patients. For *cis*-eQTL mapping, SNPs were mapped to all genes within 500 kilobases. 3,602,220 possible *cis*-eQTL associations were tested using the conventional approach of regressing gene expression level against genotype, using the following linear regression model (fit for each SNP-gene pair):

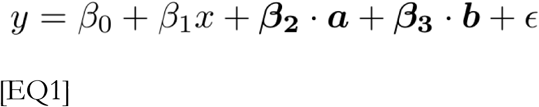

Where *y* is the gene expression value; *x* is the genotype encoded as 0, 1 or 2; *a* is the 3 genotype principal components used to estimate ancestry; *b* is the 35 PEER factors and *ε* is the residual error term. For each model, a P-value for the eQTL was calculated by a t-test on the *β_1_* term.

### Identifying cancer eQTLs using a linear model with an interaction term (the interaction model)

The model to identify cancer eQTLs is similar to the model described above but also includes tumor purity, calculated by the CPE[15] method, as a covariate and a term for the interaction of tumor purity and genotype. The model is derived in “Supplementary Information — Model Derivation”. The model is as follows:

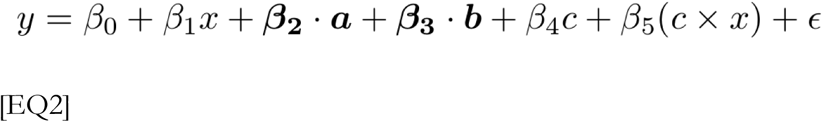

The terms are as in EQ1, but with the addition of *c*, which represents the CPE estimate of tumor purity (0 < *c* < 1) and *c* × x the interaction of tumor purity and genotype. Critically, tumor purity is encoded such that 0 represents 100% cancer cells and 1 represents 100% normal cells, meaning that the *β*_1_ term will have extrapolated an effect size at 100% cancer cells. As above, the P-value for each eQTL was calculated by a t-test on the *β*_1_ term. A similar model to EQ2 was proposed by Westra et al.[17], who successfully used it to test for eQTLs mediated by cell type proportions by testing an interaction term (*β*_5_ in EQ2). Our application to cancer relies on the following methodological innovations: In Westra et al. PC1 of their gene expression data was used a proxy for cell type proportion (term *c* in EQ2, but not bounded by 0 and 1); here, we use actual estimates of the cell type proportion, bounded by 0 and 1—in this case the proportion of tumor-associated normal cells. The consequence of this is that the main effect *β*_1_ now represents an extrapolated estimate of the eQTL effect size at 0% tumor-associated normal cells, equivalent to 100% cancer cells. Thus, we recover cancer eQTLs by testing this main effect, rather than the interaction term, which is actually a measure of how the magnitude of an eQTL differs between the two cell types (as previously described in Westra et al.). Note 1: We also fit these models with gene copy number and methylation status included as a covariate (for Supplementary Figure 3 and Supplementary Tables 4 & 5). Note 2: in EQ2 bold typeface represents vectors and the 35 PEER factors were re-estimated accounting for the tumor purity covariate not included in EQ1.

### Comparing eQTL profiles between breast cancer cells (TCGA) and normal breast tissue (GTEx)

GTEx V6 summary data, including effect sizes and associated standard errors for each SNP-gene pair, were obtained from the GTEx Portal. Cancer eQTL effects (*β*_1_ in EQ2) were compared for a given SNP-gene pair between TCGA and GTEx using the effect size and associated standard error in each dataset. A Z-score for the difference between these effects was calculated as follows[36,37]:

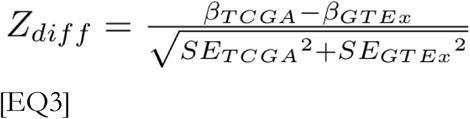

The *SE* terms refer to the standard error estimates associated with the eQTL effect *(β_TCGA_* and *β_GTEx_*) in TCGA and GTEx respectively. P-values were calculated from these Z-scores using the probability density function for a normal distribution.

### Gene set analysis of differential eQTLs between TCGA and GTEx using GOseq

Gene set analysis, which was used to identify differentially enriched biological processes between GTEx and TCGA eQTLs, was performed using the GOseq[38] package in R. We considered a gene differentially regulated if it had at least one associated eQTL that was significantly different (calculated using EQ3, *FDR* < 0.05) between TCGA breast cancer and GTEx breast tissue. All genes expressed in both TCGA breast cancer and GTEx normal breast tissue were used as the background list. The GOseq approach allowed us to use a six-knot monotonic spline function to control for the increased probability of a gene appearing in the foreground list (i.e. differentially regulated), given an increased number of associated SNPs. GOseq has previously been applied to control for similar confounders in RNA-seq[38] and methylation[39] analysis.

### Imputation of TCGA SNP data

We used the Michigan Imputation Sever v1.0.0[40] to impute genotypes for TCGA patients for the breast cancer GWAS risk variants that were not directly genotyped on the Affymetrix SNP 6.0 array used by TCGA. We used the Haplotype Reference Consortium (HRC version r1.1 2016)[41] reference panel. In addition to initial genotype calling and quality control (QC) done by TCGA, we QC’ed germline genotypes further by removing SNPs with MAF < 0.05, SNPs with missing genotype call rate >0.02, patients with missing call rate >0.02 and Hardy-Weinberg Equilibrium (HWE) *P* < 1 × 10^−6^ using Plink[42]. The input data were further validated and QC’ed by the server, followed by pre-phasing with Eagle v2.3[41] and imputation with Minimac3[43].

### METABRIC breast cancer data

The METABRIC data were obtained with permission from the European Genotype Archive. Raw Affymetrix Genome-Wide Human SNP Array 6.0 CEL files were obtained from archive EGAD00010000164. We called genotypes from these files using the Birdseed v2 algorithm under the default configuration implemented in Affymetrix Genotyping Console. Notably, these data were measured from tumor tissue and are thus less reliable than genotypes called from blood (as in TCGA); however, the METABRIC authors have previously used these genotypes for eQTL mapping, and demonstrated that the results were reasonably consistent with those obtained from genotypes generated from matched normal tissue[22]. We filtered SNPs with >.05 missing genotypes, MAF <0.05 and only retained SNPs also included in the final TCGA analysis. The METABRIC “discovery” (n = 997) normalized gene expression data were obtained from archive EGAD00010000210. We retained genes also included in the TCGA analysis and mapped each gene to a normal distribution with mean 0 and standard deviation 1. Covariates for expression heterogeneity and population structure were estimated and SNPs were mapped to genes as in the TCGA analysis above. Note that the PEER algorithm did not converge on our METABRIC expression dataset, thus we estimated expression heterogeneity using principal component analysis, applied to an expression dataset where other model covariates (population structure, purity) had been regressed out. CPE tumor purity estimates cannot be created in METABRIC as the required data types are not all available in this cohort; thus, we approximated CPE tumor purity in METABRIC by fitting a lasso regression model to CPE tumor purity estimates and gene expression in the TCGA cohort, then applied this model to METABRIC expression data; for consistency we also mapped these estimates to the same quantiles of the TCGA CPE data. Similarly to TCGA, eQTLs were then mapped using the “conventional” and “interaction” models in EQ1 & EQ2.

### Figures and data analysis

All data analysis was performed using R. Figures were created using the base plotting functions or the ggplot2 package. Because of the non-standard eQTL mapping pipeline, conventional eQTL mapping tools were not used, thus the models were fit using the *lm()* function in R. All false discovery rates were estimated using the Benjamini and Hochberg method. Most of the data analysis was performed using the Bionimbus Protected Data Cloud[44].

## Declarations

### Availability of data and materials

The code to reproduce the results in this paper are on GitHub: https://github.com/paulgeeleher/cancerEqtls and Open Science Framework: https://osf.io/z7uyp/

Note that permission must be obtained from TCGA and METABRIC to obtain access to germline genotype information.

### Ethics approval and consent to participate

N/A

### Author contributions

PG and RSH conceived the study. PG developed the statistical methods, performed the analysis and drafted the paper. CS provided support/insight in statistical methods, deconvolution approaches and analysis. JF performed exploratory initial analysis. AN and ANB provided analytical support. FW and ZZ performed genotype imputations. RG and ZZ provided computational resources and support. RSH supervised the study. All authors approved/edited the final manuscript.

## Acknowledgments

The Genotype-Tissue Expression (GTEx) Project was supported by the Common Fund of the Office of the Director of the National Institutes of Health, and by NCI, NHGRI, NHLBI, NIDA, NIMH, and NINDS. We acknowledge the Genomics Data Commons and Bionimbus Protected Data Cloud for data acquisition and analysis services and the patients and research groups who generated the TCGA data. This study also makes use of data generated by the Molecular Taxonomy of Breast Cancer International Consortium (METABRIC), funding for which was provided by Cancer Research UK and the British Columbia Cancer Agency Branch.

